# Varapp: A reactive web-application for variants filtering

**DOI:** 10.1101/060806

**Authors:** Julien Delafontaine, Alexandre Masselot, Robin Liechti, Dmitry Kuznetsov, Ioannis Xenarios, Sylvain Pradervand

## Abstract

**Summary:** Varapp is an open-source web application to filter variants from large sets of exome data stored in a relational database. Varapp offers a reactive graphical user interface, very fast data processing, security and facility to save, reproduce and share results. Typically, a few seconds suffice to apply non-trivial filters to a set of half a million variants and extract a handful of potential clinically relevant targets. Varapp implements different scenarios for Mendelian diseases (dominant, recessive, de novo, X-linked, and compound heterozygous), and allows searching for variants in genes or chromosomal regions of interest.

**Availability:** The application is made of a Javascript front-end and a Python back-end. Its source code is hosted at https://github.com/varapp. A demo version isavailable at https://varapp-demo.vital-it.ch. The full documentation can be found at https://varapp-demo.vital-it.ch/docs.

## 1 Introduction

Whole exome and gene panel sequencing have become the methods of choice to identify disease causing genetic variants. There is a current effort towards standardization of sequencing data processing for variants identification (Gargis et al, 2015), and the variant call format (VCF) has become the de-facto standard for representing variants and genotypes information. In order to allow flexible and reproducible data exploration and interpretation, VCF fileswith genome annotations can be integrated in a relational database using the Gemini softwarepackage (Paila et al., 2013). However, bioinformatics skills are required to interrogate command-line tools such as Gemini.

Here, we present Varapp, a reactive graphical user interface application to filter genetic variants that could be seen as a companion open-source software to the Gemini platform. The key features of Varapp are: (1) Privacy: no upload to a cloud or external server; (2) Reproducibility: keep track of the annotation/versions and filtering parameters used to produce the results; (3) Ease of use: fast filtering based on family pedigree, including complex scenarios such as compound heterozygous; (4) Reactivity: apply a filter in one click and see the result immediately; (5) Easy sharing of the results: send a URL to a colleague, and he sees the current state of your analysis just as you do.

## 2 Implementation

The server side of Varapp is a RESTful web service written in Python. It reads the various Gemini databases available for a given authenticated user and returns the requested variants in JSON format. For each new filters combination, it also calculates the number of variants passing each of the available filters. It uses a low-level bit masking strategy to filter by genotypes (implemented in a C extension) and a Redis cache (http://redis.io) to obtain fast response times. In the worst case presented in Table 1, it took < 2.5 seconds to returnthe results froma dataset of 438,963 variants and a compound heterozygous disease scenario.Varapp response time scales linearly with the total number of variants and the number of samples in the dataset.Preliminary tests with a whole genome dataset of 5,235,000 variants show a response time of 8.3 seconds for the dominant scenario. The gain over the standard Gemini API depends onthenumber of variants filtered in the SQL table before decompressing the genotype information. With a small database, simple disease inheritance model and stringent criteria, Gemini API performs as well as Varapp, but it can be 100x slower in other situations. Varapp performances can be further improved by parallelization.

The web interface is a Javascript application using the React framework (https://facebook.github.io/react/).It queries a Python backend to display the filtered variants in a lazy table,and provides a set of Table 1. Variant filtering benchmarks (in seconds)intuitive one-click filters that refresh the table upon every user action. It also allows tocustomize the samples selection, choose the relevant annotations, exportthe result to different formats, and save any state of the research using a system of bookmarks. The URL alwaysmirrors the application’s state so that it can be shared with other people with asimple copy-and-paste.

**Table I.** Varapp and Gemini API were used to interrogate variant databases of different size with different parameters: no disease scenario, dominant or compound het inheritance, quality filter=PASS, max frequency in any ExAC, ESP or 1000 genomes=1%. Computer is a MacBookPro with Intel Core i7, 2 cores 3.1Ghz, 16GB RAM. Values are means of 10 independent requests.

| API | Variant number in database | No scenario [s] | Qual+Freq [s] | Dominant [s] | Dominant Qual+Freq [s] | Compound het [s] | Compound het Qual+Freq [s] |
| --- | --- | --- | --- | --- | --- | --- | --- |
| Varapp | 66,082 | 0.320 | 0.316 | 0.265 | 0.457 | 0.464 | 0.338 |
| Varapp | 175,525 | 0.529 | 0.552 | 0.544 | 1.186 | 1.178 | 0.909 |
| Varapp | 438,963 | 1.276 | 1.342 | 1.158 | 2.428 | 2.474 | 2.169 |
| Gemini | 66,082 | 4.9231 | 0.3645 | 2.9453 | 0.3163 | 11.4827 | 1.0211 |
| Gemini | 175,525 | 10.8971 | 0.895 | 7.3737 | 0.7623 | 27.2766 | 1.6076 |
| Gemini | 438,963 | 30.0354 | 5.6588 | 19.7737 | 5.0928 | 322.3148 | 21.6362 |

**Fig. 1.**
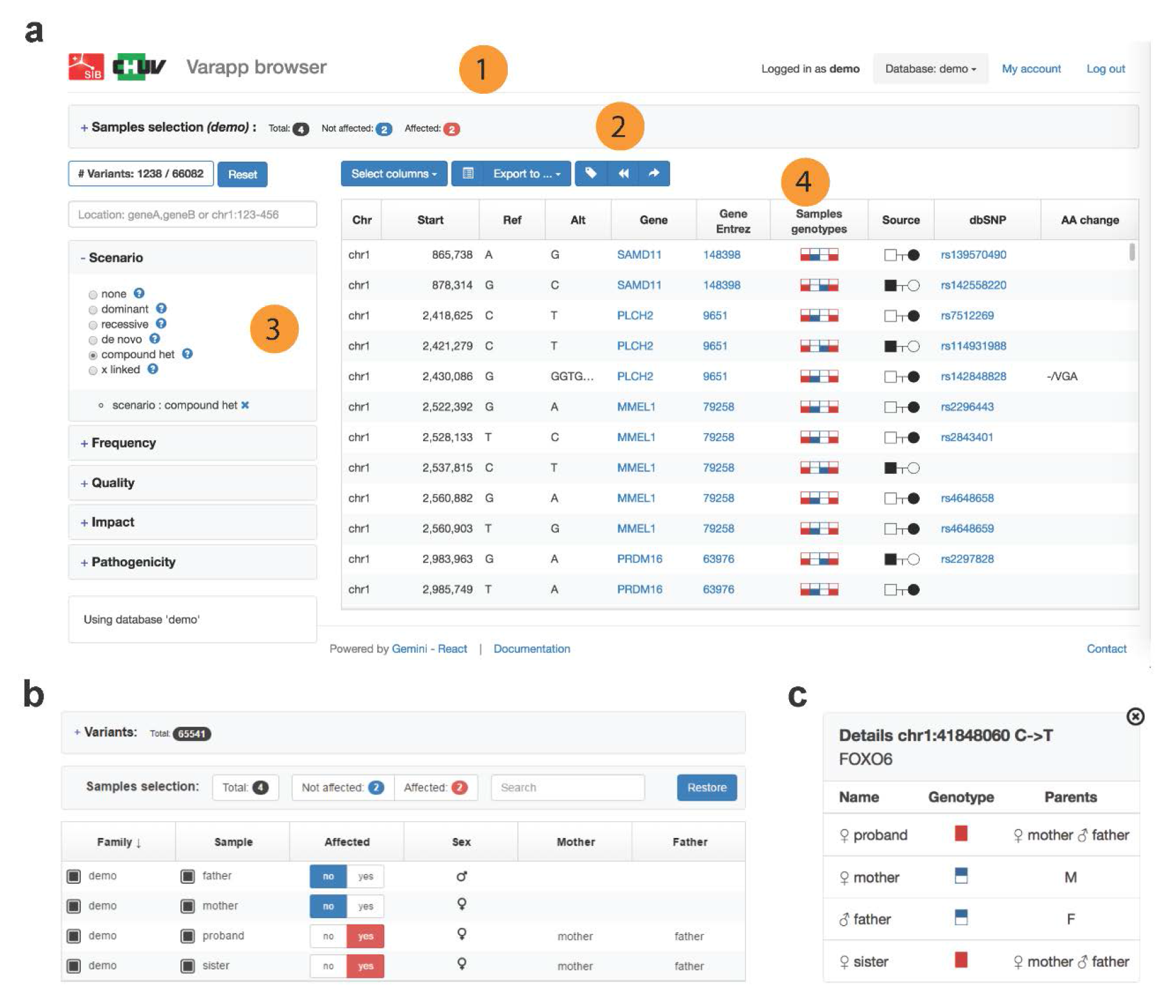
Varapp user interface. (a) Variant visualization panel is made of different sections. (1) Header with data set selection and account management. (2) Samples selection summary. Clicking on this section opens the sample selection window. (3) Filters grouped by categories and location. (4) Variants table. (b) Sample selection panel with pedigree information.(c) Lookup window. Different actions will trigger a new window with details about the clicked item. For instance, a genotypes box shows details about samples containing the variant.

The documentation provides a detailed description of data preparation, database loading and application deployment. Varapp can be installed on a local computer using the Django lightweight development server, or on a production web-server using Apache and WSGI (the Python standard for web servers and applications). For the web interface, a pre-compiled distribution is available.

## 3 Use cases

Varapp helps researchers identify causative variants in two different approaches: (1) familial idiopathic diseases (typically addressed by exome sequencing), and (2) diseases with known contributing genes (typically addressed by gene panel sequencing). In the first case, we consider all variants of one family and search for the variants that segregate with the disease. Varapp provides filters for dominant, recessive, de novo, X-linked, and compound heterozygous mode of inheritance. In the second case, we search for variants in a list of genes or genomic intervals and identify patient carriers for mutations potentially pathogenic. In both cases, the lists of variants can be refined using additional filters for (i) population frequencies, (ii) variant quality metrics, (iii) genome location and impact on protein sequence, and (iv) inference on protein function (Fig. 1a).Family and/or separate samples are selected through the ‘Samples’ panel (Fig. 1b). Disease status can be modified in the user interface for the purpose of filtering without updating the back-end database. Varapp provides a look-up window that helps identifying samples bearing a particular variant, especially useful with large cohorts of patients (Fig. 1c). The variants table can be exported in tabular text or VCF format. Varapp also generates analysis reports containing the selected samples, the filtering parameters and the annotation sources and versions.

## 4 Discussion

Varapp is aimed at biomedical research laboratories that want to centralize their different variant datasets from different projects inside a common querying system with project-based control access and possibility to share data and analysis results between collaborators. In such a system data can stay inside the institution where they were produced; there is no need to upload them to a cloud or an external server. This can be of importance regarding the legal framework to which the genetic data are bound to. Indeed, data protection directives regulates transborder data flows in many countries. Therefore, VCF upload to variant assessment software located outside the country (or the institution) can be prohibited.

Varapp is not an annotation tool (it relies on Gemini and VEP to annotate the input VCF),but it provides a graphical user interface to query relational databases containing pedigrees, variants and annotations information. There are open source tools to filter variants based on population frequency, quality or genomic location (Shameer et al. 2015). However, tools like Annovar (Wang et al. 2010), KGGSeq (Li et al. 2012), exomeSuite (Maranhao et al. 2014) or VariantMaster (Santoni et al. 2014) are command-line programs that cannot be easily used by investigators without bioinformatics expertise. There is a web-server version of Annovar (Chang and Wang 2012), but variant filtering isnot reactive in the sense that annotation/filtering jobs are submitted to a queuing system and user has to wait for a response email from the server. VariantDB is another web-based application, but it requires to set up complex filtering strategies (Vandeweyer et al. 2014). VCF-miner is GUI-based and was built to accept any arbitrary annotations from VCF files (Hart et al. 2016). It is not bound to a particular annotation tool and accepts any species, but it lacks predefined scenarios for familial disease and does not offer easy iterations over filtering parameters.

Another advantage of Varapp is the comfortable navigation from samples to variants and vice-versa, making it particularly adequate to work with cohorts and identify patients with mutations in genes of interest. To summarize, Varapp was built with user-friendliness and direct user-experience feedback in mind and possesses the unique advantages of a simple navigation, a minimal number of clicks to obtain significant results, and a very fast response time.

## Acknowledgements

We thank Norine Voisin, Lucie Gueneau, Katrin Mannik and Berit Kolk for providing guidance and use-cases during the application development and Alexandre Reymond for his lasting support to innovative technology develop-ment simplifying analytical workflows.

